# Improved metagenomic analysis with Kraken 2

**DOI:** 10.1101/762302

**Authors:** Derrick E. Wood, Jennifer Lu, Ben Langmead

## Abstract

Although Kraken’s *k*-mer-based approach provides fast taxonomic classification of metagenomic sequence data, its large memory requirements can be limiting for some applications. Kraken 2 improves upon Kraken 1 by reducing memory usage by 85%, allowing greater amounts of reference genomic data to be used, while maintaining high accuracy and increasing speed five-fold. Kraken 2 also introduces a translated search mode, providing increased sensitivity in viral metagenomics analysis.

---

Assigning taxonomic labels to sequencing reads is an important part of many computational genomics pipelines for metagenomics projects. Recent years have seen several approaches to accomplish this task in a time-efficient manner^1–3^. Kraken^4^ used a memory-intensive algorithm that associates short genomic substrings (*k*-mers) with lowest common ancestor (LCA) taxa. Kraken and related tools like KrakenUniq^5^ have proven highly efficient and accurate in other tool comparisons^6,7^. But Kraken’s high memory requirements force many researchers to either use a reduced-sensitivity MiniKraken database^8,9^, or to build and use many indexes over subsets of the reference sequences^10,11^. Its memory requirements can easily exceed 100 GB^7^, especially when the reference data includes large eukaryotic genomes^12,13^. Here we introduce Kraken 2, which provides a major reduction in memory usage as well as faster classification, a spaced-seed searching scheme, a translated search mode for matching in amino acid space, and continued compatibility with the Bracken^14^ species-level quantification algorithm.

Kraken 2 addresses the issue of large memory requirements through two changes to Kraken 1’s data structures and algorithms. While Kraken 1 used a sorted list of *k*-mer/LCA pairs indexed by minimizers^15^, Kraken 2 introduces a probabilistic, compact hash table to map minimizers to LCAs. This table uses one-third of the memory of a standard hash table, at the cost of some specificity and accuracy. Additionally, Kraken 2 only stores minimizers (of length ℓ, ℓ ≤ *k*) from the reference sequence library in the data structure, whereas Kraken 1’s stored all *k*-mers. Kraken 2’s index for a reference database consisting of 9.1 Gbp of genomic sequence uses 10.6 gigabytes of memory at classification time. Kraken 1’s index for the same reference set uses 72.4 gigabytes of memory for classification (**Figure 1a, Supplementary Table S1**). In general, a Kraken 2 database is about 15% as large as a Kraken 1 database over the same references (**Supplementary Figure S1**).

**Figure 1.**
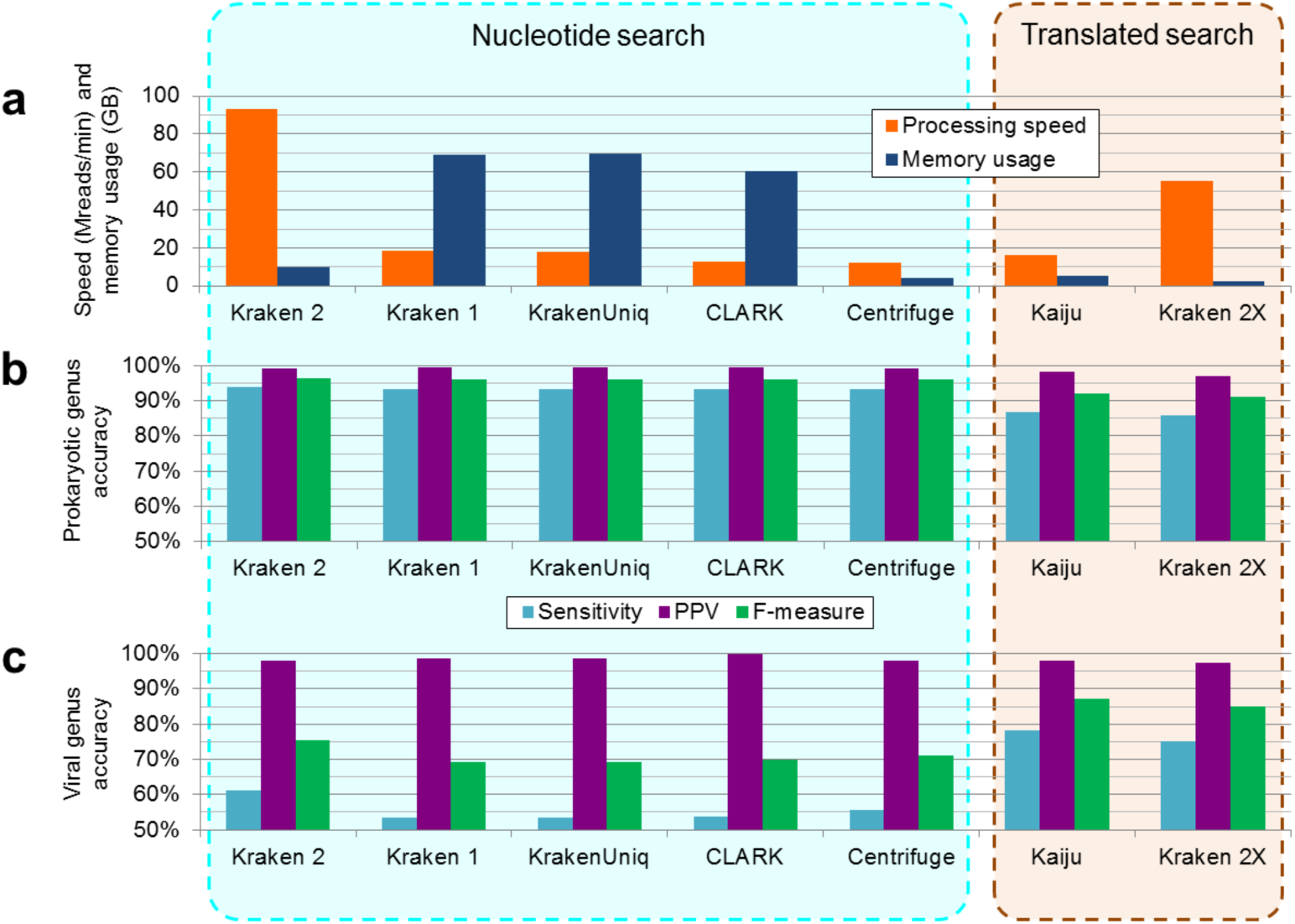
Comparison between Kraken 2 and other tools. **(a)** Processing speed (in millions of reads per minute) and memory usage (measured by maximum resident set size, in gigabytes) are shown for each classifier, as evaluated on 50 million paired-end simulated reads with 16 threads. Accuracy results are shown for **(b)** 40 prokaryotic genomes and **(c)** 10 viral genomes. Results here are shown for sensitivity, positive predictive value (PPV), and F-measure as evaluated on a per-read basis at the genus rank, with 1000 reads simulated from each genome. The strains from which reads were simulated were excluded from the reference libraries for each classification tool. “Kraken 2X” is Kraken 2 using translated search against a protein database. Full results for these strain-exclusion experiments are available in **Supplementary Table S1**.

Kraken 2’s approach is faster than Kraken 1’s because only the distinct minimizers from the query (read) trigger accesses to the hash table. A similar minimizer-based approach has proven useful in accelerating read alignment^16^. Kraken 2 additionally provides a hash-based filtering approach that subsamples the set of minimizer/LCA pairs included in the table, allowing the user to specify a target hash table size; smaller hash tables yield lower memory footprint and higher classification throughput at the expense of lower classification accuracy (**Supplementary Table S2**).

Kraken 2 also features other improvements to accuracy and runtime. A new translated search mode (“Kraken 2X”) uses a reduced amino-acid alphabet and increases sensitivity on viral datasets compared to nucleotide-based search. Block- and batch-based parsing within the critical section is used to improve thread scaling, in a manner similar to that used in recent versions of Bowtie 2^17^. We also added a form of spaced seed search and automated masking of low-complexity reference sequences to improve accuracy.

To assess the accuracy and performance of Kraken 2, we selected 40 prokaryotic and 10 viral genomes for which we had reference genomes for at least 2 sister sub-species and at least 2 sister species (**Supplementary Table S3**). We then created a reference genome (or protein) set that excluded the 50 taxa for the genomes we selected. This reference set and taxonomy were held constant between the various classifiers we examined, avoiding any confounding due to differences in reference database. A similar approach has been recently used for this same purpose in another study^7^.

We simulated one million Illumina 100 x 100 nt paired-end reads from each of the 50 selected genomes, for a total of 50 million reads (25 million fragments). We processed these data with four nucleotide search-based sequence classification programs (Centrifuge^1^, CLARK^2^, Kraken 1^4^, and KrakenUniq^5^), and a translated search classifier (Kaiju^3^). We additionally processed these data with Kraken 2, using several different databases created with different parameters (**Methods**).

This strain-exclusion approach mimics the real-world scenario where reads likely originate from strains that are genetically distinct from those in the database. The addition of simulated sequencing errors also provides further genetic distance between the test data and the reference sequences. Through this approach, we sought to avoid overly optimistic estimates of a classifier’s performance.

We found that Kraken 2 exhibited similar, and often superior, per-sequence accuracy to the other nucleotide classifiers, and that Kraken 2X provided similar (though slightly lower) accuracy compared to Kaiju (**Figure 1b, Supplementary Table S1**). The nucleotide-based classifiers exhibited lower accuracy on the viral read data than did the translated-search classifiers, demonstrating the advantage of translated search in scenarios marked by high genetic variability and sparsity of available reference genomes^3^.

We then examined the runtime and memory requirements of each program. Kraken 2 provided substantial increases in processing speed, classifying paired-end data at over 93 million reads per minute while using 16 threads, a speed over 5 times faster than Kraken 1, the next-fastest classifier (**Figure 1a, Supplementary Table S1**). Additionally, Kraken 2 exhibited superior thread scaling to Kraken 1 (**Supplementary Table S4**). Kraken 2’s memory requirement is also 15% of Kraken 1’s, and only 2.5 times as much as that of the least memory-intensive classifier we examined, Centrifuge. With respect to the translated search programs, Kraken 2X is over 3 times faster and uses 47% less memory than Kaiju.

To determine if Kraken 2 exhibited similar analytical performance on real sequencing data, we classified read data from the FDA-ARGOS project^18^. We compared the fragment classifications obtained by the various classification programs to the taxonomic labels attached to the corresponding ARGOS experiment. Kraken 2 exhibits similar genus-level concordance and discordance statistics to the other nucleotide-search classifiers, while Kraken 2X exhibits similar but less agreement with the ARGOS labels than does Kaiju (**Supplementary Table S5**). These results agree with those obtained in the strain-exclusion experiment on simulated data.

As a continuation of the strain-exclusion experiments, we applied Bracken^14^ to the Kraken 1 and Kraken 2 results, estimating species and genus-level sequence abundance for prokaryotic species. Bracken uses a Bayesian algorithm to integrate reads Kraken classified at higher taxonomic levels into the abundance estimates. Although the true strain-level taxa are excluded from the database, Bracken recaptured most of the true genus-level and species-level sequence abundances using both Kraken 2 and Kraken 1 classification results. Comparing the results, the Bracken estimates were more accurate with Kraken 2 than with Kraken 1 at both the genus and species level, likely owing to Kraken 2’s higher sensitivity (**Figure 2**).

**Figure 2.**
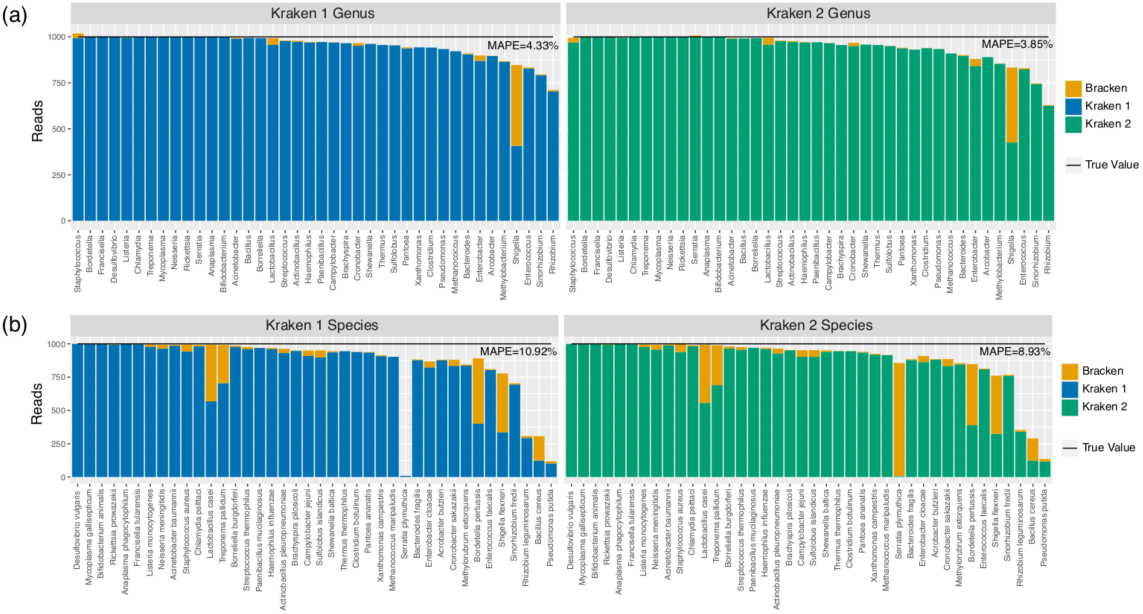
Bracken performance on strain exclusion simulated prokaryotic data. For each of the 40 genomes examined, we removed the genome from the reference genome set, and simulated 1000 paired-end reads from the genome. We used Kraken 1 and Kraken 2 to build databases with the same reference set, and to classify each simulated fragment. We then used Bracken with both programs to estimate **(a)** genus sequence abundance estimation and **(b)** species sequence abundance estimation. “MAPE” is mean absolute percentage error, as defined in the **Methods**.

As databases of assembled genomes continue to grow, databases of reference sequences used for metagenomics studies will also grow^19,20^. We presented Kraken 2, an extremely memory-efficient metagenomics classification tool that replaces Kraken 1’s *k*-mer database with a probabilistic data structure that is substantially smaller, allowing 6-7 times as much reference data compared to Kraken 1. The algorithms introduced in Kraken 2 to subsample the set of genomic substrings also provide Kraken 2 with the ability to further reduce the size of its database and accelerate the processing of sequencing data. We showed Kraken 2’s accuracy is comparable to that of Kraken 1 and other competing tools, consistent with other studies^6,7^. We also showed that its new translated search mode has accuracy approaching that of the protein-focused Kaiju tool, while using less memory and runtime. Also, Kraken 2 is compatible with the Bracken software for species-level quantification, making Kraken 2 straightforwardly usable for that application.

In the future it will be important to consider additional use cases for Kraken 2. For example, other data structures similar to our compact hash table, such as the counting quotient filter^21^, could be implemented and used in computing environments and applications that may benefit from a particular data structure’s design and properties. Additionally, the KrakenUniq^5^ tool uses the HyperLogLog sketch^22^ to estimate the number of distinct *k*-mers matched at each node of the taxonomy, a statistic that is used in turn to better determine presence-or-absence of individual genomes. We plan to add this functionality in the future, as it enables applications in diagnosis of infections where the infectious agent is present at low abundance.

## Methods

### Compact hash table

The hash table used by Kraken 2 to store minimizer/LCA key-value pairs is very similar to a traditional hash table that would use linear probing for collision resolution, with some modifications. Kraken 2’s compact hash table (CHT) uses a fixed-size array of 32-bit hash cells to store key-value pairs. Within a cell, the number of bits used to store the value of the key-value pair will vary depending on the number of bits needed to represent all unique taxonomy ID numbers found in the reference sequence library; this was 17 bits with the standard Kraken 2 database in September 2018. The value is stored in the least significant bits of the hash cell, and must be a positive integer. Values of zero represent empty cells. Within the remaining bits of the hash cell, the most significant bits of the key’s hash code (a compact hash code) are stored. Searching for a key *K* in the CHT is done by computing the hash code of the key *h*(*K*), then linearly scanning the table array starting at position *h*(*K*) mod |*T*| (where |*T*| is the number of cells in the array) for a matching key. Examples of this search process – including both key/value insertion and querying – are shown in **Supplementary Figure S2.** In Kraken 2, the hash function *h* used is the finalization function from MurmurHash3^23^.

Compacting hash codes in this way allows Kraken 2 to use 32 bits for a key-value pair, a reduction compared to the 96 bits used by Kraken 1 (64 bits for key, 32 for value). But it also creates a new way in which keys can “collide,” in turn impacting the accuracy of queries to the CHT. Two distinct keys can be treated as identical by the CHT if they share the same compact hash code and their starting search positions are close enough to cause a linear probe to encounter a stored matching compact hash code before an empty cell is found. This property gives the CHT its probabilistic nature, in that two types of false positive query results are possible: either (a) a key that was not inserted can be reported as present in the table, or (b) the values of two keys can be confused with each other. In Kraken 2, the former error is indeed a false positive, whereas the latter results in a less specific LCA being assigned to the minimizer (**Supplementary Figure S2**). The probability of either of these errors is <1% with Kraken 2’s default load factor of 70% (**Supplementary Figure S3**). The adverse effect on read-level classification is further mitigated by the algorithm Kraken 2 uses to combine information from across the read, which is unchanged from Kraken 1 and utilizes information from all *k*-mers in a sequence to counteract low-frequency erroneous LCA values that could be returned by a key-value store.

The probabilistic nature and comparisons involving parts of a key’s hash code make the CHT similar to the counting quotient filter (CQF) described by Pandey *et al*.^21^ Like the CQF, Kraken 2’s CHT features high locality of memory access during an individual query due to the linear probing that the CHT employs. Unlike the CQF, however, our CHT does not allow the full hash code to be recovered from a stored value (the CQF’s *remainder*), and so we are unable to resize a CHT once it is instantiated. Additionally, our CHT has an additional possibility of error compared to the CQF, where two keys that do not have the same full hash code but share a truncated hash code will be treated as identical. The CQF can avoid such “soft” hash collisions.

### Internal taxonomy of a Kraken 2 database

While Kraken 1 used the taxonomy provided by the user without modification, Kraken 2 makes some modifications to its internal representation of the taxonomy that cause that representation to differ from the user-provided taxonomy. First, Kraken 2 finds a minimal set of nodes in the user-provided taxonomy. This minimal set consists of all nodes to which a reference sequence is assigned, as well as all of those nodes’ ancestors; vertices between nodes in this set remain as they were in the user-provided taxonomy, maintaining the tree structure in the internal representation. Kraken 2 then assigns nodes in the minimal set sequentially increasing internal taxonomy ID numbers using a breadth-first search (BFS) beginning at the root, with the root having an internal ID number of 1. This BFS provides a guarantee that ancestor nodes will have smaller internal ID numbers than their descendants; an example of this numbering is shown in **Supplementary Figure S2**. Kraken 2 stores a mapping of its internal taxonomy numbers to the external taxonomy ID numbers to make its results more easily interpretable, and performs all output using the external taxonomy ID numbers.

Kraken 2’s use of this internal taxonomy representation allows for easier computation of the LCA of two nodes because the ID numbers themselves give information as to their relative depths in the tree, while the National Center for Biotechnology Information (NCBI) taxonomy IDs lack this property. The internal taxonomy representation also allows Kraken 2 to use the minimal number of bits for storage of taxonomy ID numbers, giving maximal space for the compact hash codes and reducing the probability of CHT errors (or “hash table collisions”, as we describe elsewhere in this paper).

A Kraken 2 database consists of a CHT and this internal taxonomy representation. Typical databases will be built using the NCBI taxonomy^24^ but users can override this default to create custom databases for atypical use cases.

### Minimizer-based subsampling

In contrast to Kraken 1’s use of all *k*-mers in the standard use case, Kraken 2 subsamples the set of genomic substrings and inserts only the distinct minimizers into its database. We define the ℓ bp minimizer of a *k*-mer (ℓ ≤ *k*) to be the lexicographically smallest canonical ℓ-mer found within the *k*-mer. An ℓ-mer is called canonical if it is lexicographically less than or equal to its reverse complement. Note that if *k*=ℓ, no subsampling occurs and Kraken 2 inserts the same substrings into its data structure that Kraken 1 would. The default values for Kraken 2, *k*=35 and ℓ=31, were determined after analysis of the parameter sweep results we show in **Supplementary Table S2**.

Kraken 2 determines which ℓ-mers are minimizers by use of a sliding window minimum algorithm, in contrast to Kraken 1’s implementation which examined each *k-*mer anew. This allows for a faster determination of minimizers, as less work is required when moving from one *k-*mer to the next overlapping *k-*mer (in terms of computational complexity, the new approach uses an average of *O*(1) time to calculate a new minimizer vs. Θ(*k*) time with the older algorithm). The sliding window minimum calculation uses a double-ended queue (or “deque”) in which canonicalized candidate ℓ-mers are inserted in the back, along with the candidates’ position in the original sequence. As a new candidate is encountered, enqueued candidates are removed from the back of the deque until the candidate at the back has a greater value than the new candidate (as determined by lexicographical ordering). The new candidate is then pushed onto the back of the deque.

Once a *k-*mer’s worth of ℓ-mers have been processed in this way, the front of the deque contains the minimizer of that *k-*mer. This property is then maintained during scanning subsequent bases by removing the front element in the deque if it is from a position in the original sequence that is not in the current *k-*mer. In this way, the front element of the deque holds the minimizer of the *k-*mer currently being examined.

We further augmented the sliding window algorithm to include the exclusive-or (XOR) shuffling operation from Kraken 1. This operation serves to permute the ordering of the ℓ-mers when calculating minimizers, and helps to avoid a bias toward low-complexity ℓ-mers when selecting the minimizer of a *k*-mer^4,15^. To shuffle, we calculate the XOR value of the ℓ-mer and a predefined constant, and use this value as the “candidate” that is put in the deque. When the original ℓ-mer value is needed again, the operation is reversed by XORing a second time with the same constant.

### Spaced seed usage

Spaced *k*-mers, a similar concept to spaced seeds, have been shown to improve the ability to classify reads within the Kraken framework^25^. Kraken 2 uses a simple spaced seed approach where a user specifies an integer *s* when building a database that indicates how many positions in the minimizer will be masked (*i.e.*, not considered when searching). Beginning with the next-to-rightmost position in the minimizer, every other position is masked until *s* positions have been masked. For example, if *s*=3 and ℓ =12, the positions in the bit-string 1111 1101 0101 with a “0” would be masked. Kraken 2’s default value for *s* is 7 and was determined after analysis of the parameter sweep results we show in **Supplementary Table S2**.

The canonical ℓ-mers that are minimizer candidates are masked with the spaced seed mask prior to their insertion into the deque for the sliding window calculation. By performing canonicalization of the minimizer candidates prior to applying the spaced seed mask, we ensure the result is the same whether applied to the ℓ-mer or its reverse complement.

### Hash-based subsampling

Kraken 2 estimates the required capacity of the hash table given the *k*, ℓ, and *s* values chosen along with the sequence data in a database’s reference genomic library. Some users will not have access to large memory computers, and therefore this estimate may be greater than the maximum possible hash table size that they can work with. To aid such users, Kraken 2 allows them to specify a maximum size when building a database. If the estimated required capacity is larger than the maximum requested size, then the minimizers will be subsampled further using a hash function. Given an estimated required capacity *S*’ and a maximum user-specified capacity of *S* (*S* < *S*’), we can calculate the value *f*=*S*/*S*’, which is the fraction of available minimizers that the user will be able to hold in their database. A minimum allowable hash value of *v*=(1-*f*)·*M* can also be calculated, where *M* is the maximum value output by hash function *h*. Any minimizer in the reference library with a hash code less than *v* will not be inserted into the hash table. This value *v* is also provided to the classifier so that only minimizers with hash codes greater than or equal to *v* will be used to probe the hash table, saving the search failure runtime penalty that would be incurred by searching for minimizers guaranteed not to be in the hash table.

### Evaluation of *k*-mer level discordance rates

At a *k*-mer level, there are two main types of discordance between Kraken 1 and Kraken 2’s results: those caused by two distinct *k*-mers sharing the same minimizer (a “minimizer collision”), and those caused by two distinct minimizers being indistinguishable by the CHT (a “hash table collision”). Minimizer collisions are not always damaging. When it occurs between *k*-mers from very closely-related genomes, such a collision might detect true homology even in the face of single nucleotide polymorphisms and/or sequencing error. That said, minimizer collisions between *k*-mers from distantly-related genomes could produce either elevated LCA values (if both genomes are in the reference library) or incorrectly classified *k*-mers (if one of the genomes is not in the reference library). Hash table collisions are a consequence of the probabilistic nature of the CHT, and can also cause either elevated LCA values or incorrectly classified *k*-mers (**Supplementary Figure S2**). We note that these different discordant results are all at a *k*-mer level, and may not always affect a query sequence’s classification due to the many *k*-mers’ worth of data that are used to classify a query sequence; aside from slight modifications to handle the subsampling methods we use in Kraken 2, the classification method of Kraken 2 is identical to Kraken 1.

We wished to estimate the rate at which these collisions would cause discordance at a *k*-mer level between the Kraken 1 and Kraken 2 results. To do so, we selected a specific bacterial genome for which we had neighboring genomes at each taxonomic rank from species to phylum. The selected genome was our “reference sequence”, and eight others were progressively more taxonomically distant from the reference sequence. We list the nine genomes used in these experiments in **Supplementary Table S6**. We additionally created a synthetic genome with 4 Mbp of uniformly random DNA. Together, these ten sequences form a set of “query sequences” and are the basis for our evaluation of collision rates. For these experiments, we used the default Kraken 2 values of *k*=35, ℓ=31, and *s*=7, unless otherwise noted.

To determine the rates of discordance caused by minimizer collisions, we compared each of the ten query sequences’ *k*-mers to the set of reference sequence *k*-mers. For each sequence, the minimizer collision rate is the proportion of distinct *k*-mers in a query sequence that (a) are not in the set of reference sequence *k*-mers and (b) share a minimizer with a reference sequence *k*-mer. The various sequences’ minimizer collision rates are summarized in **Supplementary Table S7**. We hypothesized that the minimizer collision rate would be influenced by the length of the minimizer used, due to the length’s direct relationship to the number of possible minimizers. To test this, we repeated the minimizer collision rate estimation experiment focusing on the reference genome and using the random synthetic genome as the sole query sequence. Setting *k*=35 and *s*=0, we varied the ℓ parameter from 8 to 31. Minimizer lengths greater than 15 had collision rates under 1%. Minimizer lengths greater than 22 had zero collisions. Full results are shown in **Supplementary Figure S4**.

To determine the rates of discordance caused by hash table collisions, we compared each of the ten query sequences’ minimizers to a CHT populated with the reference sequence minimizers. The CHT was created with a load factor of 70% and 15 bits reserved for the truncated hash code (the same parameters used in Kraken 2’s standard database in September 2018). For each sequence, the hash table collision rate is the proportion of distinct minimizers in a query sequence that (a) are not minimizers in the set of reference sequence minimizers and (b) are reported by the CHT as being inserted in the hash table. The various sequences’ hash table collision rates are summarized in **Supplementary Table S8**. To investigate the impact of load factor and truncated hash code size on hash table collision rates, we repeated the hash table collision rate experiment, but focused only on the reference genome and used the random synthetic genome as the sole query sequence. We used the same default values of *k*, ℓ, and *s* as before (35, 31, and 7, respectively), and calculated hash table collision rates while varying both the load factor and truncated hash code size. The impact of these two parameters on hash table collision rates is shown in **Supplementary Figure S3**. The parameters adopted for Kraken 2’s default mode had an error rate of 0.016%, consistent with the results seen when comparing genomes of different species (**Supplementary Table S8**).

### Processing of a standard genomic reference library

The CHT’s modest memory requirements, and the additional savings yielded by minimizer-based subsampling, allow more reference genomic data to be included in Kraken 2’s standard reference library. Whereas Kraken 1’s default database had data from archaeal, bacterial, and viral genomes, Kraken 2’s default database additionally includes the GRCh38 assembly of the human genome^26^ and the “UniVec_Core” subset of the UniVec database^27^. We include these in Kraken 2’s default database to allow for easier classification of human microbiome reads and more accurate classification of reads containing vector sequences.

Additionally, we have implemented masking of low-complexity sequences from reference sequences in Kraken 2, by using the “dustmasker”^28^ (for nucleotide sequences) and “segmasker”^29^ (for protein sequences) tools from NCBI. Using the tools’ default settings, nucleotide and protein sequences are checked for low-complexity regions, and those regions identified are masked and not processed further by the Kraken 2 database building process. In this manner, we seek to reduce false positives resulting from these low-complexity sequences, similarly to the build process for Centrifuge^1^.

### Populating the Kraken 2 hash table

Kraken 2 begins building a CHT by first estimating the number of distinct minimizers present in the reference library for the selected values of *k*, ℓ, and *s*. This is done through a form of zeroth frequency moment estimation^30^ where Kraken 2 creates a small set structure implemented with a traditional hash table. In this set *Q*, we insert only the distinct minimizers that satisfy the criterion *h*(*m*) mod *F* < *E*, where *h*(*m*) is the hash code of the minimizer *m*, and *E* << *F* (in practice, Kraken 2 uses *E* = 4 and *F* = 1024). We then find the estimate of the total number of distinct minimizers by multiplying the number of satisfactory distinct minimizers (|*Q*|) by *F*/*E*. This form of estimation requires storing in memory only a fraction of all distinct minimizers (approximately *E*/*F*), and allows us to quickly set the capacity of our CHT properly without needing to first store all elements in it.

After estimating the number of distinct minimizers *D=*|*Q*|(*F*/*E*) present in the reference library, Kraken 2 then allocates memory for a CHT containing *D*/0.7 hash table cells. We selected the divisor of 0.7 so that the resultant hash table will have approximately 30% of its cells remain empty after the population of the CHT (*i.e.*, the CHT will have a load factor of 70%). As stated earlier, the cells of this table are 32 bits each, and so the total memory required for Kraken 2’s CHT is 32*D*/0.7 bits, or 4*D*/0.7 bytes.

Kraken 2 then proceeds to scan each genome in the reference library. Each genome must be associated with a taxonomic ID number so that Kraken 2 can calculate LCA values; genomes without associated taxonomy IDs are therefore not processed by Kraken 2. For a minimizer *M* in a genome G, Kraken 2 attempts to insert a key-value pair containing *M* (key) and the taxonomic ID *T* (value) associated with G into the CHT. If the CHT does not report that *M* was previously inserted, then the <*M, T*> key-value pair will be inserted, indicating that the LCA of *M* is currently *T*. If *M* was previously inserted into the CHT, with LCA value *T\**, then its associated LCA value is updated to equal the LCA of *T* and *T\**. All minimizers are processed in this way; once the reference library’s minimizers are all processed, the LCA values are properly set for each of the minimizers and the database build is complete. The LCA operation is both commutative and associative, facilitating parallel index construction.

### Classification of a sequence fragment with Kraken 2

Kraken 2 classifies sequence fragments similarly to Kraken 1, with modifications to facilitate minimizer- and hash-based subsampling. For each *k*-mer in an input sequence, Kraken 2 finds its minimizer and, if it is distinct from the previous k-mer’s minimizer, uses it as a key to probe the CHT. If the minimizer matches a key in the CHT, Kraken 2 considers the associated LCA value to be the *k*-mer’s LCA. Classification then proceeds in the same manner as Kraken 1, taking note of how many *k*-mer hits mapped to each taxa, constructing a pruned classification tree, and using the leaf of the maximally scoring root-to-leaf path of that tree to classify the sequence^4^. If hash-based subsampling was used to build the CHT, each minimizer has its hash code compared against the table’s maximum allowable hash code, and minimizers with higher-than-allowed hash codes are not searched against the CHT. Any *k*-mer containing an ambiguous nucleotide code is also not searched against the CHT.

### Parsing of input files

Previous work by Langmead *et al*.^17^ has shown the importance of removing parsing work from critical sections, *i.e.*, portions of the program that can be executed by only one thread at a time. Kraken 2 uses two different methods to defer a majority of parsing work from the critical section to thread-local execution. The first method (referred to as “batch deferred” parsing by Langmead *et al*.) reads a set number of lines (40,000 in Kraken 2) of input in a thread-local buffer within the critical section, and then parses the input within a single thread’s execution. This method is used to perform reading of paired-end FASTQ input, where the lengths of a fragment’s mates can be different and reading a consistent number of lines from both input files is necessary to ensure a thread is working with complete mate pairs. For FASTA or single-end FASTQ input, Kraken 2 instead uses a more efficient method that reads in a set number of bytes (3 MB in Kraken 2) of input into a thread-local buffer within the critical section, and continues reading input into that buffer until a record boundary is found, at which point a thread leaves the critical section and parses its input. These modifications allow Kraken 2 to more efficiently use multiple threads than did Kraken 1 (**Supplementary Table S4**).

### Translated search

To perform translated search, Kraken 2X first builds a database from a set of reference proteins in the same manner that Kraken 2 does for nucleotide sequences. The usual alphabet of 20 amino acids is reduced to 15 using the 15-character alphabet of Solis^31^; we add a single additional value representing selenocysteine, pyrrolysine, and translation termination (stop codons). This gives us 16 characters in our reduced alphabet, allowing us to represent a character with 4 bits. Minimizers of reference proteins are calculated using the same methods for nucleotide sequences (*i.e.*, using spaced seeds if requested and a sliding window minimum algorithm), but reverse complements are not calculated and by default *k*=15, ℓ=12, and *s*=0.

When searching against a protein minimizer database, Kraken 2X translates all six reading frames of the input query DNA sequences into the reduced amino acid alphabet. Minimizers from all six frames are pooled and used to query the CHT, and therefore all contribute to the Kraken 2X classification of a query sequence.

### Generation of data for strain exclusion experiments

We downloaded the reference genome and protein data used for the clade exclusion experiments from NCBI in January 2018 from the archaeal, bacterial, and viral domains. We also downloaded the taxonomy from NCBI at this same time. Using the taxonomy ID information for each sequence, we obtained a set of all taxonomy IDs represented by the reference genomes. From this set, we selected a subset that had both two sister sub-species taxa present and two sister species taxa present in the set of reference genomes. From this subset, 40 prokaryotic taxonomy IDs and 10 viral taxonomy IDs were selected arbitrarily to be the strains of origin for our experiments. The strains selected are listed in **Supplementary Table S3**.

We then created two sets of reference libraries where all data from the strains of origin were removed, one for the nucleotide sequences and one for the protein sequences. We used Mason 2^32^ to simulate 100 bp paired-end Illumina sequence data from our strains of origin, with 500,000 fragments being simulated from each strain. When simulating the reads, we used the default options for simulating sequencing errors with Mason 2’s “mason_simulator” command. These defaults caused the simulator to simulate sequencing errors at rates of 0.4% for mismatches, 0.005% for insertions, and 0.005% for deletions. We combined simulated reads from the strains of origin into a single set of read data. We also shuffled the order of the fragments in this set to control for ordering effects that might affect runtime.

### Execution of strain exclusion experiments

In brief, we used the nucleotide search-based classification programs (Kraken 1, KrakenUniq, Kraken 2, CLARK, and Centrifuge) to build a strain-exclusion database from reference genomes, and we used the translated search-based classification programs (Kraken 2X and Kaiju) to build a strain-exclusion database from reference protein sequences. We compared Kraken 2 and Kraken 2X (both using the code base from Kraken 2.0.8) against Kraken 1.1.1, KrakenUniq 0.5.6, CLARK 1.2.4, Centrifuge 1.0.3-beta, and Kaiju 1.5.0. Because CLARK requires a rank to be specified at the time of building a database, and our evaluations center on genus-rank accuracy, we built a CLARK database for the genus rank for our evaluation work in this paper.

Classifiers received the simulated read data as paired-end FASTQ input. To evaluate runtime and memory usage, we sought to eliminate the performance impact of reading or writing from disk or from a network storage location. To accomplish this, we copied simulated read data and classifier databases onto a RAM filesystem and directed the classifiers to read input from and write output to that RAM filesystem.

Accuracy was evaluated on a smaller subset of the simulated data containing 1,000 fragments per genome of origin, or 50,000 fragments in total. To obtain processing speed and memory usage information, we ran each classifier using 16 threads on 25 million sequences’ worth of simulated read data. We used the taskset command to restrict each classifier to the appropriate number of processors (*e.g.*, “taskset -c 0-15” was used with our 16 thread experiments); this ensures that a classifier that uses an external process to aid in its execution has that process’ runtime properly counted against its runtime here. The “/usr/bin/time -v” command provided us with elapsed wall clock time and maximum resident set size data (memory usage) for each experiment, and allowed us to verify that no major page faults were incurred by a classifier during its execution (the absence of which indicates minimal disk- or network-related input/output effects on the runtime). Classifiers were run on a computer with 32 Xeon 2.3 GHz CPUs (16 hyperthreaded cores) and 244 GB of RAM.

### Evaluation of accuracy in strain exclusion experiments

We evaluated the accuracy of each classifier at a per-fragment level, with respect to a particular taxonomic rank. Each fragment had a known true sub-species taxon of origin, which implied a true taxon of origin at both the species and genus ranks, which is where we measured accuracy. We now describe how we counted true positive (TP), false negative (FN), vague positive (VP), and false positive (FP) results at the genus and species levels. We describe this at the genus level specifically, but the analogous procedure was also used at the species level. For a given true genus of origin, a TP classification is a classification at that genus or at a descendant of that genus. Because we excluded the strains of origin from our reference databases, we expected all classifiers to make incorrect strain-level classifications, and so allow classifications of descendants of the true genus to be judged as TP. We define an FN classification as a failure of a classifier to assign any classification to a sequence, and a VP classification as a classification at an ancestor of the true genus of origin. Finally, we define an FP classification as a classification that is incorrect; that is, not at the true genus of origin, nor an ancestor or descendant of that true genus. These four categories are mutually exclusive, and all fragments run through a classifier will have their classification (or lack thereof) categorized by one of these categories.

These categories are different from those typically used for binary classification problems; they are used here because these methods can make classifications that are not at leaves of the taxonomic tree, but are still correct. For example, a classification of an *Escherichia coli* fragment as *Escherichia* would be evaluated as TP for genus-rank accuracy, but as VP for species-rank accuracy. Classification of that same fragment as *Vibrio* would be evaluated as FP at any rank below class (because the LCA of *Vibrio* and *Escherichia* is the class taxon Gammaproteobacteria) and would be evaluated as TP for the class rank and above.

Using these categories, we define rank-level sensitivity as the proportion of input fragments that were true positive classifications, or TP/(TP+VP+FN+FP). We define rank-level positive predictive value (PPV) as the proportion of classifications that were true positives (excluding acceptable but vague classifications), or TP/(TP+FP). Along with these definitions of rank-level sensitivity and PPV, we also define an F-measure as the harmonic mean of those two values.

### Evaluation of thread scaling efficiency

To evaluate Kraken 1’s and Kraken 2’s ability to efficiently use multiple threads, we performed an experiment using the strain exclusion databases and simulated read data we describe previously in this section. We ran both Kraken 1 and Kraken 2 on the same data using 1, 4, and 16 threads. The two programs were run once on the data as paired-end read data, and once as single-end read data. Read data and Kraken database files were all placed on a RAM filesystem and the “taskset” command was used to limit the classifier programs to only as many cores as the number of threads being used. These conditions mirror those of our main strain exclusion experiments, only varying the number of threads between the various runs of the classifiers. Results for this experiment are shown in **Supplementary Table S4**. In short, Kraken 2 exhibits superior speedup with respect to the number of threads allocated compared to Kraken 1. This is especially true for paired-end reads.

### FDA-ARGOS experimental concordance evaluation

The FDA-ARGOS (dAtabase for Reference Grade micrObial Sequences) project provides sequencing experiments for many microbial isolates^18^. We used the NCBI’s Sequence Read Archive^33^ to find all 1,392 experiments related to the FDA-ARGOS project (accession PRJNA231221). Because some tools are unable to properly process reads of differing lengths, we selected only those 263 experiments that were run on an Illumina HiSeq 4000 instrument and produced 151 bp reads. We then randomly selected one experiment from each genus to download, and used reservoir sampling to select a subset of 10,000 paired-end fragments from each selected experiment. We also removed experiments for which our strain-exclusion reference genome set did not have a reference genome of the same species as the sequenced isolate. These steps yielded 25 experiments’ worth of data, for 250,000 paired-end fragments in total. Using the strain-exclusion databases created earlier, we then used each classifier to classify the data and examined the percentage of each experiment’s fragments that were classified.

Because the FDA-ARGOS data are from real sequencing experiments, several factors could explain discordance between a classifier’s results and the experiments’ assigned taxa, including evolutionary distance between sequences and reference data, low-quality sequencing runs, and contamination. The true causes of such discordance may not be discernable, and even when they are, they often require an in-depth examination of the sequencing and reference data. For these reasons, we do not report sensitivity and PPV for these data because we cannot be certain of the true taxonomic origin of each individual fragment of real sequencing data. Rather, we evaluated the concordance of the SRA-assigned taxa with the fragments’ classifications at the genus rank, and report for each classifier the following quantities: (a) the percentage of fragments with a concordant classification at the genus rank; (b) the percentage of fragments with a discordant classification at the genus rank; (c) the percentage of fragments with a classification of an ancestor of the SRA-assigned genus taxon; and (d) the percentage of fragments that were not classified. The results of this concordance evaluation are provided in full in **Supplementary Table S5**.

### Parameter sweeps

We examined various values for parameters to ensure Kraken 2’s default parameters would provide an advantageous balance of accuracy, classification speed, and memory usage. Specifically, we looked at parameters relating to minimizer-based subsampling (*k* and ℓ), hash-based subsampling (*f*=*S*/*S’*), and spaced-seed usage (*s*). For Kraken 2, we performed two parameter sweeps, with one focused on minimizer-based subsampling, and one focused on hash-based subsampling. The first parameter sweep looked at values for ℓ in the interval [25, 31], values for k in the interval [ℓ, ℓ+10], and values for s in the interval [0, 7]; the second parameter sweep looked at values of ℓ in the interval [25, 31], fixed *k*= ℓ, values for *f* in the set {0.125, 0.25, 0.5}, and values for *s* in the interval [0, 7]. We also performed a third parameter sweep, focused on translated search (Kraken 2X), where we looked at values for ℓ in the interval [11, 15], values for *k* in the interval [ℓ, ℓ+3], and values for *s* in the interval [0, 3].

Each parameter sweep used the strain exclusion data that we previously created to build databases, and we used the same accuracy and timing methods for these databases that we did in the cross-classifier comparison. The results of the first two parameter sweeps, run on nucleotide databases, are provided in **Supplementary Table S2**, while the results of the third parameter sweep, run on protein databases, are provided in **Supplementary Table S9**. We note that the parameter sweeps yielded a large number of parameter combinations giving approximately the same, near-optimal levels of accuracy. This suggests performance is not overly sensitive to particular parameter settings.

### Evaluation of database sizes of Kraken 1 and Kraken 2

We began by shuffling the reference DNA sequences in our strain exclusion set, and recorded the total number of bases in each sequence. We modified Kraken 2’s capacity estimator to report an estimate of the number of distinct minimizers after each sequence processed, rather than only after all sequences are processed. Finally, we ran the capacity estimator twice on the shuffled genomic data, once with *k*=31, ℓ=31, *s*=0 (corresponding to Kraken 1’s defaults – effectively counting the number of distinct *k*-mers); and again with *k*=35, ℓ=31, *s*=7 (Kraken 2’s defaults).

The size of a Kraken 1 database is a function of the number of distinct *k*-mers in the reference data. If there are *X* distinct *k*-mers, the size of Kraken 1’s database.kdb (sorted list of *k*-mer/LCA pairs) file will be 1072 + 12*X* bytes; the 1072 byte term is the size of the Jellyfish/Kraken header data, and 12 bytes are used for each *k*-mer/LCA pair. The database.idx (minimizer offset index) file is 8589934608 bytes, a function of Kraken 1’s default minimizer length of 15. The full database size is the sum of sizes of those two files.

Similarly, the size of a Kraken 2 hash table is a function of the estimate of the number of distinct minimizers in the reference data. If there are an estimated *Y* distinct minimizers, Kraken 2’s hash table will be 32 + ⌊4*Y*/0.7⌋ bytes in size (representing 32 bytes of metadata, and using 4 bytes per cell and a load factor of 0.7).We used the estimates of the numbers of distinct *k*-mers and distinct minimizers to calculate the database sizes of Kraken 1 and Kraken 2 for successively larger subsets of the strain exclusion set. The results of this evaluation are shown in **Supplementary Figure S1**, with raw data available in **Supplementary Table S10**.

Reviewing the results when all genomic sequences were added, our results indicate that the number of distinct *k*-mers is approximately 3.1 times the number of distinct minimizers for the settings we have selected for Kraken 1 and Kraken 2. It is not possible to draw a direct relationship between the number of distinct *k*-mers or minimizers and the number of sequence bases processed. For example, homology between similar strains and species will cause number of distinct *k*-mers/minimizers to grow slower than the total number of bases. Examining the linear-term coefficients from the database-size expressions (12*X* and 4*Y*/0.7) indicates a Kraken 2 database will be approximately 15% of the size of a Kraken 1 database of the same reference data; this is because *X* ≈ 3.1*Y*, and (4/0.7)/(12*3.1) = 0.15. When we examine the full reference set, the 15% estimate is consistent with the ratio of Kraken 2’s hash table size (10.456 GB) to Kraken 1’s database.kdb file size (77.490 – 8.589 = 68.901 GB), which is 10.456/68.901=0.152.

### Bracken experiments on strain exclusion data

We first generated Bracken metadata from each of the Kraken 1 and Kraken 2 reference libraries used in the strain exclusion experiments. We then used Bracken to estimate genus- and species-level abundance from the Kraken 1 and Kraken 2 classification results on the prokaryotic strain exclusion read data. Due to the low sequence similarity between our simulated viral reads and the strain-exclusion reference data, none of the nucleotide-search programs exhibited high sensitivity on these reads, including Kraken 1 and Kraken 2. Such low classification rates prevent Bracken from inferring taxonomy for a large proportion of the viral reads. Additionally, the taxonomy for viruses has several examples where species are not grouped by ancestry, and lack similarity in both gene organization and genomic sequence^34^. For these reasons, we chose to exclude the simulated viral reads from our analysis of Bracken.

For overall evaluation of the accuracy of Bracken in these strain exclusion experiments, we calculated the mean absolute percentage error (MAPE):

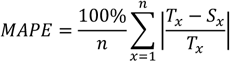

where *S*_*x*_ is the estimated number of reads and *T*_*x*_ is the true number of reads for taxon *x*. In this strain exclusion experiment, *n* = 40, the total number of distinct prokaryotic species and genera in the sample and *T*_*x*_ = 1000 for each taxon.

## Data and source code availability

We have made the data for our strain exclusion experiments publicly available for download, including all reference sequences, taxonomy, and simulated read data^35^. Code to generate all databases from these reference sequences, to generate simulated read data, and to run the comparison of classifiers’ accuracy and performance is also available for public download^36^.

Kraken 2’s source code is open-access, licensed under the MIT license, and available in a GitHub repository^37^.

## Acknowledgements

The authors would like to thank James R. White and Steven Salzberg for helpful discussions about the manuscript. BL and DEW were supported by NSF grant IIS-1349906. BL was additionally supported by NIH grant R01-GM118568. JL was supported by NIH grant R35-GM130151.

## Author Contributions

DEW and BL designed the algorithms for Kraken 2. DEW developed the Kraken 2 software. DEW, JL, and BL designed the experiments. DEW and JL performed experiments. DEW, JL, and BL prepared and reviewed the manuscript.

## References

1. Kim, D., Song, L., Breitwieser, F. P. & Salzberg, S. L. Centrifuge: rapid and sensitive classification of metagenomic sequences. Genome Res 26, 1721–1729 (2016).

2. Ounit, R., Wanamaker, S., Close, T. J. & Lonardi, S. CLARK: fast and accurate classification of metagenomic and genomic sequences using discriminative k-mers. BMC Genomics 16, 236 (2015).

3. Menzel, P., Ng, K. L. & Krogh, A. Fast and sensitive taxonomic classification for metagenomics with Kaiju. Nat Commun 7, 11257 (2016).

4. Wood, D. E. & Salzberg, S. L. Kraken: ultrafast metagenomic sequence classification using exact alignments. Genome Biol 15, R46 (2014).

5. Breitwieser, F. P., Baker, D. N. & Salzberg, S. L. KrakenUniq: confident and fast metagenomics classification using unique k-mer counts. Genome Biol 19, 198 (2018).

6. Lindgreen, S., Adair, K. L. & Gardner, P. P. An evaluation of the accuracy and speed of metagenome analysis tools. Sci Rep 6, 19233 (2016).

7. Ye, S. H., Siddle, K. J., Park, D. J. & Sabeti, P. C. Benchmarking Metagenomics Tools for Taxonomic Classification. Cell 178, 779–794 (2019).

8. Eyice, Ö. et al. SIP metagenomics identifies uncultivated Methylophilaceae as dimethylsulphide degrading bacteria in soil and lake sediment. ISME J 9, 2336 (2015).

9. Merelli, I. et al. Low-power portable devices for metagenomics analysis: Fog computing makes bioinformatics ready for the Internet of Things. Futur Gener Comput Syst 88, 467–478 (2018).

10. Lu, J. & Salzberg, S. L. Removing contaminants from databases of draft genomes. PLOS Comput Biol 14, e1006277 (2018).

11. Donovan, P. D., Gonzalez, G., Higgins, D. G., Butler, G. & Ito, K. Identification of fungi in shotgun metagenomics datasets. PLoS One 13, e0192898 (2018).

12. Meiser, A., Otte, J., Schmitt, I. & Grande, F. D. Sequencing genomes from mixed DNA samples - evaluating the metagenome skimming approach in lichenized fungi. Sci Rep 7, 14881 (2017).

13. Knutson, T. P., Velayudhan, B. T. & Marthaler, D. G. A porcine enterovirus G associated with enteric disease contains a novel papain-like cysteine protease. J Gen Virol 98, 1305–1310 (2017).

14. Lu, J., Breitwieser, F. P., Thielen, P. & Salzberg, S. L. Bracken: estimating species abundance in metagenomics data. PeerJ Comput Sci 3, e104 (2017).

15. Roberts, M., Hayes, W., Hunt, B., Mount, S. & Yorke, J. Reducing storage requirements for biological sequence comparison. Bioinformatics 20, 3363–3369 (2004).

16. Li, H. Minimap2: pairwise alignment for nucleotide sequences. Bioinformatics 34, 3094–3100 (2018).

17. Langmead, B., Wilks, C., Antonescu, V. & Charles, R. Scaling read aligners to hundreds of threads on general-purpose processors. Bioinformatics bty648 (2018).

18. Sichtig, H. et al. FDA-ARGOS: A Public Quality-Controlled Genome Database Resource for Infectious Disease Sequencing Diagnostics and Regulatory Science Research. bioRxiv 482059 (2018). doi:10.1101/482059

19. Stewart, R. D. et al. Assembly of 913 microbial genomes from metagenomic sequencing of the cow rumen. Nat Commun 9, 870 (2018).

20. Parks, D. H. et al. A standardized bacterial taxonomy based on genome phylogeny substantially revises the tree of life. Nat Biotechnol 36, 996 (2018).

21. Pandey, P., Bender, M. A., Johnson, R. & Patro, R. A General-Purpose Counting Filter: Making Every Bit Count. in Proc 2017 ACM Int Conf Manag Data 775–787 (2017). doi:10.1145/3035918.3035963

22. Flajolet, P., Fusy, É., Gandouet, O. & Meunier, F. Hyperloglog: The analysis of a near-optimal cardinality estimation algorithm. Discret Math Theor Comput Sci Proc 127–146 (2007).

23. Appleby, A. SMHasher GitHub repository. at <https://github.com/aappleby/smhasher>

24. Federhen, S. The NCBI Taxonomy database. Nucleic Acids Res 40, D136–D143 (2011).

25. Br̆inda, K., Sykulski, M. & Kucherov, G. Spaced seeds improve k-mer-based metagenomic classification. Bioinformatics 31, 3584–3592 (2015).

26. Church, D. M. et al. Extending reference assembly models. Genome Biol 16, 13 (2015).

27. The UniVec Database. at <https://www.ncbi.nlm.nih.gov/tools/vecscreen/univec/>

28. Morgulis, A., Gertz, E. M., Schäffer, A. A. & Agarwala, R. A Fast and Symmetric DUST Implementation to Mask Low-Complexity DNA Sequences. J Comput Biol 13, 1028–1040 (2006).

29. Wootton, J. C. & Federhen, S. Analysis of compositionally biased regions in sequence databases. Methods Enzymol 266, 554–571 (1996).

30. Flajolet, P. & Martin, G. N. Probabilistic counting algorithms for data base applications. J Comput Syst Sci 31, 182–209 (1985).

31. Solis, A. D. Amino acid alphabet reduction preserves fold information contained in contact interactions in proteins. Proteins Struct Funct Bioinforma 83, 2198–2216 (2015).

32. Holtgrewe, M. Mason - A Read Simulator for Second Generation Sequencing Data. Technical Report TR–B–10–06 (2010).

33. Kodama, Y. et al. The sequence read archive: explosive growth of sequencing data. Nucleic Acids Res 40, D54–D56 (2011).

34. Lawrence, J. G., Hatfull, G. F. & Hendrix, R. W. Imbroglios of Viral Taxonomy: Genetic Exchange and Failings of Phenetic Approaches. J Bacteriol 184, 4891 LP – 4905 (2002).

35. Wood, D. E. Kraken 2 Manuscript Data. doi:10.5281/zenodo.3365797

36. Wood, D. E. Kraken 2 Experiment GitHub repository. at <https://github.com/DerrickWood/kraken2-experiment-code>

37. Wood, D. E. Kraken 2 GitHub repository. at <https://github.com/DerrickWood/kraken2>

