## Supplementary Figures for "Improved metagenomic analysis with Kraken 2"

for

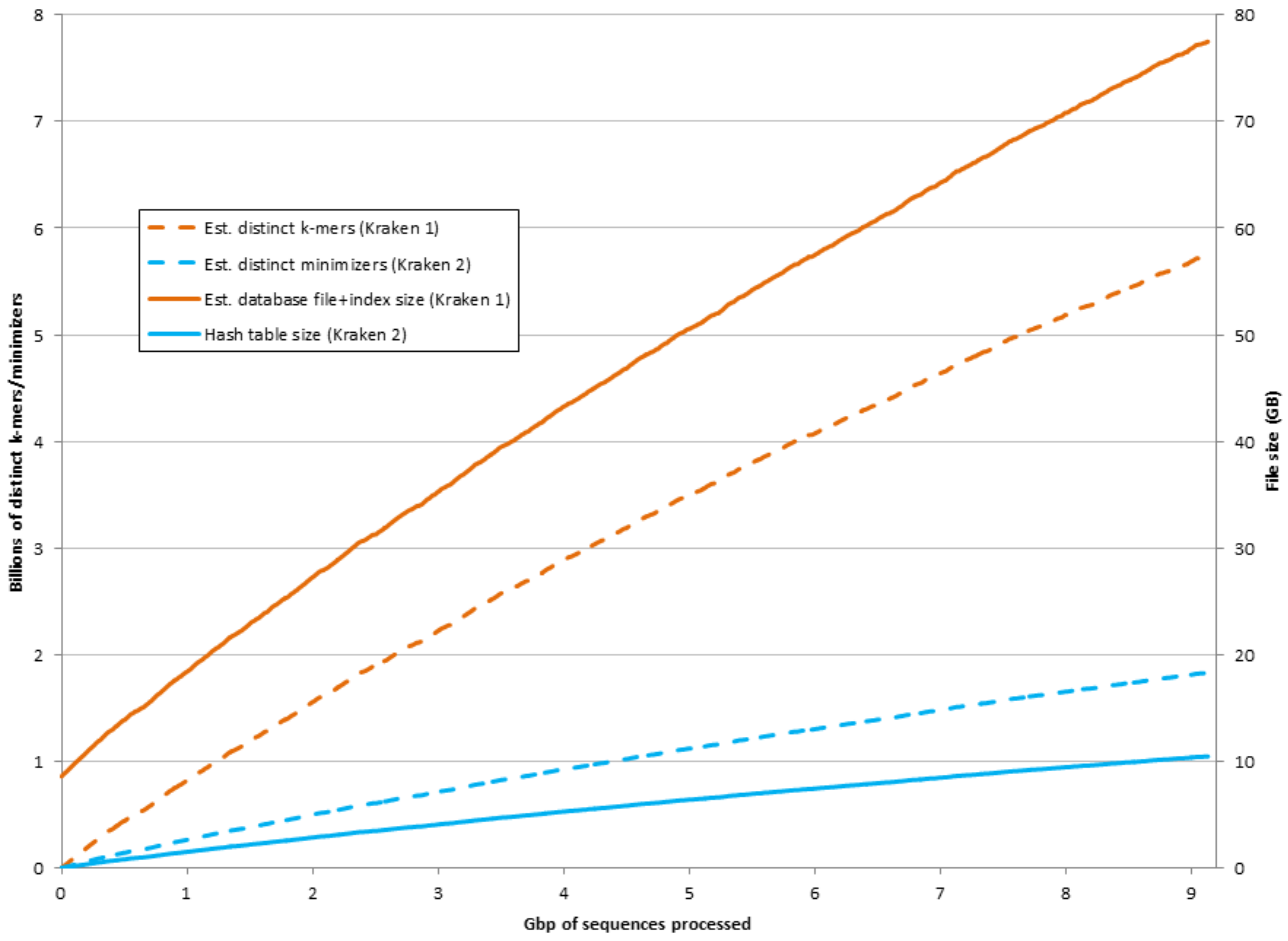


**Supplementary Figure S1. Estimation of database sizes for Kraken 1 and Kraken 2 as sequences are added to the reference set.** Over a shuffled set of the nucleotide sequences in our strain exclusion reference, we calculated progressively larger estimates of the number of distinct *k*-mers and minimizers. Because database sizes for Kraken 1 and Kraken 2 are functions of the numbers of distinct *k*-mers and minimizers, respectively, we also calculated the estimated database sizes for Kraken 1 and Kraken 2.

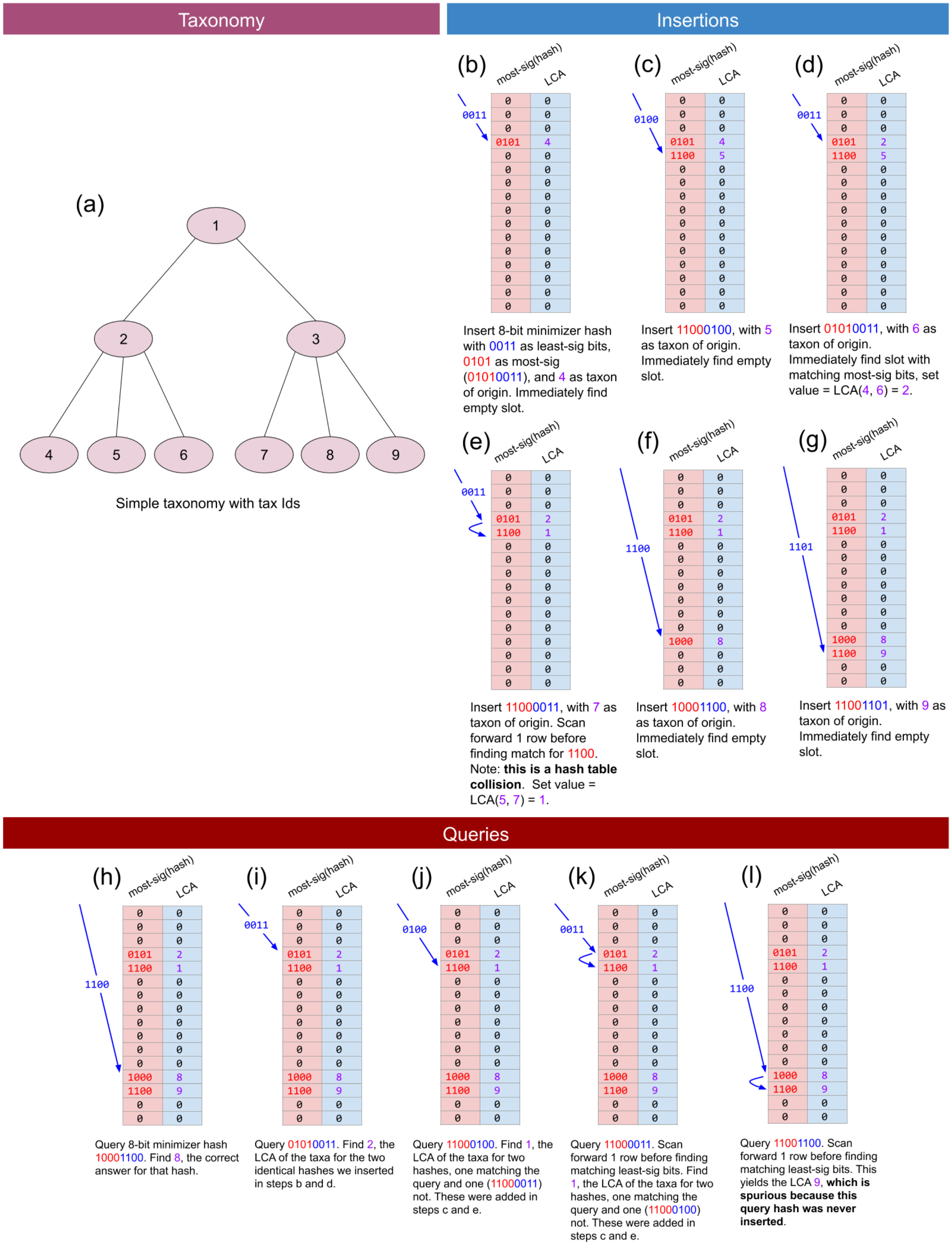


**Supplementary Figure S2. Examples of compact hash table usage with Kraken 2. (a)** An example of Kraken 2's reduced internal representation of the taxonomy with sequential ID numbering via breadth-first search. **(b)-(g)** Sequential examples of Kraken 2's insertion of minimizer/LCA pairs into a compact hash table. **(h)-(l)** Sequential examples of Kraken 2's querying of a compact hash table with a minimizer. In practice, the table and the lengths of the keys are greater than shown here. Also, since the actual table does not generally have a number of rows equal to a power of 2, we use the modulo operation to obtain the search offset (blue) rather than simply taking bits from the hash value.

| Truncated Hash Code Storage Bits | Load Factor 10% | 20% | 30% | 40% | 50% | 60% | 70% | 80% | 90% |
|---|---|---|---|---|---|---|---|---|---|
| 26 | 0.000% | 0.000% | 0.000% | 0.000% | 0.000% | 0.000% | 0.000% | 0.000% | 0.000% |
| 25 | 0.000% | 0.000% | 0.000% | 0.000% | 0.000% | 0.000% | 0.000% | 0.000% | 0.000% |
| 24 | 0.000% | 0.000% | 0.000% | 0.000% | 0.000% | 0.000% | 0.000% | 0.000% | 0.000% |
| 23 | 0.000% | 0.000% | 0.000% | 0.000% | 0.000% | 0.000% | 0.000% | 0.000% | 0.000% |
| 22 | 0.000% | 0.000% | 0.000% | 0.000% | 0.000% | 0.000% | 0.000% | 0.000% | 0.001% |
| 21 | 0.000% | 0.000% | 0.000% | 0.000% | 0.000% | 0.000% | 0.000% | 0.001% | 0.002% |
| 20 | 0.000% | 0.000% | 0.000% | 0.000% | 0.000% | 0.000% | 0.000% | 0.001% | 0.005% |
| 19 | 0.000% | 0.000% | 0.000% | 0.000% | 0.000% | 0.001% | 0.001% | 0.002% | 0.008% |
| 18 | 0.000% | 0.000% | 0.000% | 0.000% | 0.001% | 0.001% | 0.001% | 0.005% | 0.017% |
| 17 | 0.000% | 0.000% | 0.000% | 0.000% | 0.001% | 0.002% | 0.004% | 0.009% | 0.037% |
| 16 | 0.000% | 0.000% | 0.001% | 0.001% | 0.002% | 0.004% | 0.008% | 0.017% | 0.074% |
| 15 | 0.000% | 0.001% | 0.002% | 0.003% | 0.005% | 0.009% | 0.016% | 0.036% | 0.150% |
| 14 | 0.000% | 0.002% | 0.004% | 0.006% | 0.009% | 0.017% | 0.032% | 0.073% | 0.296% |
| 13 | 0.001% | 0.003% | 0.007% | 0.012% | 0.019% | 0.032% | 0.063% | 0.145% | 0.587% |
| 12 | 0.002% | 0.006% | 0.012% | 0.022% | 0.038% | 0.067% | 0.123% | 0.286% | 1.145% |
| 11 | 0.005% | 0.013% | 0.025% | 0.044% | 0.074% | 0.129% | 0.245% | 0.555% | 2.207% |
| 10 | 0.012% | 0.026% | 0.049% | 0.086% | 0.150% | 0.260% | 0.480% | 1.101% | 4.030% |
| 9 | 0.022% | 0.052% | 0.100% | 0.174% | 0.297% | 0.505% | 0.941% | 2.132% | 6.860% |
| 8 | 0.045% | 0.105% | 0.195% | 0.345% | 0.574% | 0.992% | 1.820% | 3.966% | 10.895% |
| 7 | 0.091% | 0.216% | 0.397% | 0.686% | 1.124% | 1.931% | 3.448% | 6.936% | 15.927% |
| 6 | 0.182% | 0.430% | 0.783% | 1.341% | 2.183% | 3.647% | 6.216% | 11.358% | 21.765% |

**Supplementary Figure S3. Evaluation of compact hash table error rates as a function of two variables.** The error rates of Kraken 2's compact hash table are a function of the load factor and the number of bits used to store the truncated hash code. Error rates were determined by inserting the minimizers from *P. aeruginosa* UCBPP PA14 and querying with minimizers from randomly generated sequence. Kraken 2's default database settings used 15 bits to store the truncated hash code and a load factor of 70%, which is highlighted with a red border in the figure.

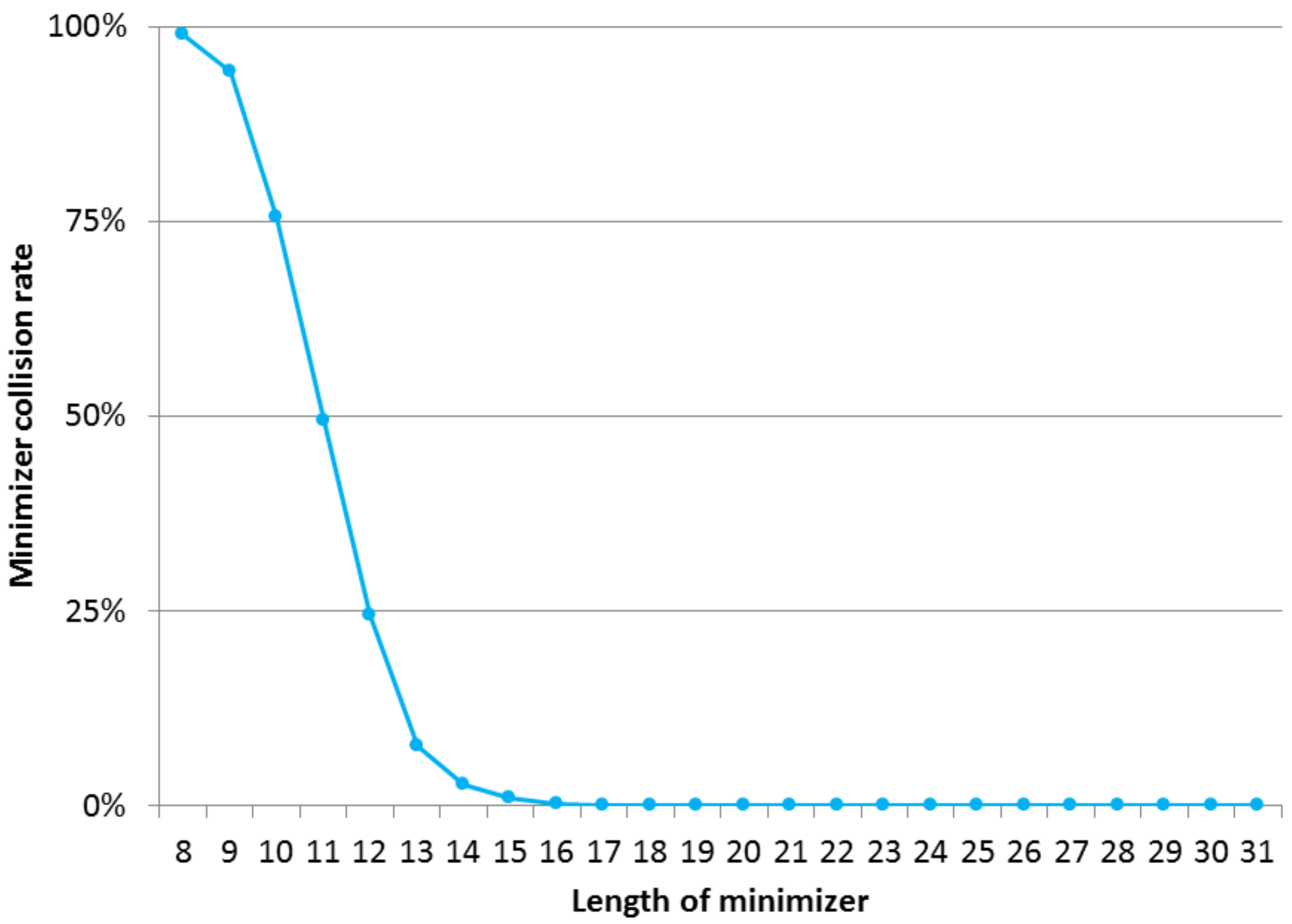


**Supplementary Figure S4. Evaluation of minimizer collision rates as a function of minimizer length.** We examined the rate at which minimizers of 35-mers from *P. aeruginosa* UCBPP PA14 would be found as minimizers of 35-mers in randomly generated sequence. Minimizer lengths ℓ were varied from 8 to 31, with no spaced minimizers used. This demonstrates the significantly lower collision rates that occur by use of the long (ℓ=31) minimizers in Kraken 2 versus use of shorter (ℓ <16) minimizers.
